# Genome-wide association study of alcohol consumption and genetic overlap with other health-related traits in UK Biobank (N=112,117)

**DOI:** 10.1101/116707

**Authors:** Toni-Kim Clarke, Mark J. Adams, Gail Davies, David M. Howard, Lynsey S. Hall, Sandosh Padmanabhan, Alison D. Murray, Blair H. Smith, Archie Campbell, Caroline Hayward, David J. Porteous, Ian J. Deary, Andrew M. McIntosh

## Abstract

Alcohol consumption has been linked to over 200 diseases and is responsible for over 5% of the global disease burden. Well known genetic variants in alcohol metabolizing genes, e.g. *ALDH2, ADH1B,* are strongly associated with alcohol consumption but have limited impact in European populations where they are found at low frequency. We performed a genome-wide association study (GWAS) of self-reported alcohol consumption in 112,117 individuals in the UK Biobank (UKB) sample of white British individuals. We report significant genome-wide associations at 8 independent loci. These include SNPs in alcohol metabolizing genes (*ADH1B/ADH1C/ADH5*) and 2 loci in *KLB,* a gene recently associated with alcohol consumption. We also identify SNPs at novel loci including *GCKR, PXDN, CADM2* and *TNFRSF11A.* Gene-based analyses found significant associations with genes implicated in the neurobiology of substance use (*CRHR1, DRD2*), and genes previously associated with alcohol consumption (*AUTS2*). GCTA-GREML analyses found a significant SNP-based heritability of self-reported alcohol consumption of 13% (S.E.=0.01). Sex-specific analyses found largely overlapping GWAS loci and the genetic correlation between male and female alcohol consumption was 0.73 (S.E.=0.09, p-value = 1.37 x 10^−16^). Using LD score regression, genetic overlap was found between alcohol consumption and schizophrenia (rG=0.13, S.E=0.04), HDL cholesterol (rG=0.21, S.E=0.05), smoking (rG=0.49, S.E=0.06) and various anthropometric traits (e.g. Overweight, rG=-0.19, S.E.=0.05). This study replicates the association between alcohol consumption and alcohol metabolizing genes and *KLB*, and identifies 4 novel gene associations that should be the focus of future studies investigating the neurobiology of alcohol consumption.

## Introduction

In 2012, 5.9% of all global deaths were attributable to alcohol and roughly a quarter of all deaths in the 20-39 years age group^1^. Over 200 diseases are linked to alcohol consumption and the proportion of the global disease burden, measured in disability adjusted life years (DALYs), is over 5%^1^. Almost a quarter of DALYs attributable to alcohol consumption were the result of neuropsychiatric disorders such as alcohol use disorders (AUD) and major depressive disorder^2^.

There is a substantial genetic component to the variation in alcohol consumption although studies of alcohol use have largely focused on AUD to date. A recent meta-analysis of twin and adoption studies estimated the heritability of AUD to be 0.49 [95%C.I. 0.43-0.53]^3^. A study of twin pairs with genome-wide SNP data estimated that 33% (S.E.=0.12) of the variance in AUD is attributable to common SNPs and SNPs in linkage disequilibrium (LD) with them^4^. Studies of the heritability of *alcohol consumption* are far fewer in number, but a study of 2877 twin pairs estimated the heritability of alcohol consumption to be 0.43 (95% C.I. 0.31-0.56). Using a sample of un-related parents, the same authors estimated ~18% of the variance in alcohol consumption to be attributable to common SNPs^5^.

Specific genetic variants have been linked to variation in alcohol consumption, the most influential being rs671 in the aldehyde dehydrogenase (*ALDH2*) gene and a cluster of variants spanning the alcohol dehydrogenase genes (*ADH1B, ADH1C, ADH5, ADH6, ADH7*) located on chromosome 4q. The catalytically inactive version of *ALDH2* encoded by the A allele of rs671 leads to slower metabolism of acetaldehyde which causes the alcohol flush reaction. This causes alcohol to be aversive for the A carriers of rs671^6,7^ As such, this polymorphism is highly protective against alcoholism in Asian populations,^8^ although it has limited impact in European and African populations where it is often monomorphic^9^. rs1229984 in *ADH1B* is also associated with an alcohol-flush reaction in Asian populations and thus protects against high alcohol consumption^10^. It is also found to associate with drinking patterns phenotypes in European and African populations where the minor allele frequency is ~1-4% ^9, 11, 12^

Genome-wide association studies (GWAS) of alcohol consumption have found few consistently replicable loci outside of the alcohol-metabolizing genes. A GWAS of alcohol consumption comprising 47,501 individuals of European ancestry found rs6943555 in *AUTS2* to be associated with alcohol consumption^13^. This locus did not replicate in a more recent GWAS of alcohol consumption in >105,000 European individuals; however, a SNP in *KLB* was associated with alcohol consumption in humans and the gene product, β-klotho, found to regulate alcohol preference in mice^14^.

Alcohol consumption has been linked to psychiatric disorders and other health related traits; however, this has generally been limited to epidemiological observations^15, 16^ or by assessing genetic overlap using twin studies^17, 18^. Levels of alcohol consumption within a population are strongly linked to cardiovascular disease, liver cirrhosis and cancer^16^. Furthermore, twin studies have shown that genetic factors overlap with alcoholism and depression^19^, ADHD and externalizing disorders^17^. Studies which have assessed the SNP-based genetic overlap between alcohol consumption and other traits have been limited to other substance abuse phenotypes. They have found that the polygenic architecture underlying alcohol consumption is shared with tobacco, caffeine and cannabis use^20, 21^.

In the present study we perform the largest GWAS of self-reported alcohol consumption in 112,117 individuals of European ancestry from the UK Biobank. We also estimate the SNP-based heritability of alcohol consumption and perform sex-specific analyses to investigate whether the phenotypic differences in alcohol consumption in males and females have a genetic basis. Furthermore, as alcohol consumption is often associated with psychiatric and other diseases and health related traits, we examined the genetic overlap between alcohol consumption and over 200 disease and behavioural traits. Using a polygenic risk score (PRS) approach we also analysed the amount of variance in self-reported alcohol consumption in a completely independent sample, Generation Scotland: the Scottish Family Health Study (GS) (N=19,858).

## Methods

### Samples

*UK Biobank:* UK Biobank (UKB) is a population based sample comprising 502,000 individuals resident in the United Kingdom between the age of 40 and 69 years^22^. During the recruitment period from 2006-2010 individuals were recruited from 22 centres across the UK to reflect a broad socioeconomic demographic and mixture of urban and rural residents. During a self-completed touchscreen interview taken at baseline appointment participants were asked about their current drinking status (never, previous, current, prefer not to say) and were asked to report their average weekly and monthly alcohol consumption of a range of drink types (red wine, white wine, champagne, spirits, beer/cider, fortified wine). These questions were accompanied by pictures providing an example of a single measure of each drink type. From these measures we derived an average intake of alcohol consumption in units per week. We excluded all former drinkers from the analysis and this left 112,117 individuals with data on both alcohol consumption and genome wide genotype data. We also repeated our analyses with never drinkers excluded (N=108,309) and found the GWAS results to be consistent with the GWAS of the full sample.

*Generation Scotland: the Scottish Family Health Study:* Generation Scotland (GS) is a family and population based study comprising 24,096 individuals aged between 18 and 99 years^23, 24^ Genome-wide genotype data is available for 19,858 individuals^25^. Alcohol consumption was assessed using a pre-clinical questionnaire. Participants self-identified as current drinkers, former drinkers or never drinkers. Average consumption was self-reported units of alcohol consumed in the previous week and these questions were prompted by a table containing the average units contained in a single measure of various drink types to assist participants in the calculation of weekly intake. Any GS individuals or relatives of GS individuals were removed from the UK Biobank sample prior to GWAS analysis.

This study obtained informed consent from all participants and was conducted under generic approval from the National Health Service (NHS) National Research Ethics Service (approval letter dated 17 June 2011, Ref 11/NW/0382) and under UK Biobank approval 4844 ‘Stratifying Resilience and Depression Longitudinally’ (principal investigator AMM). All components of GS have received ethical approval from the NHS Tayside Committee on Medical Research Ethics (REC Reference Number: 05/S1401/89) and written consent for the use of data was obtained from all participants.

### Genotyping and Imputation

*UKB:* The majority of the UKB sample was genotyped using the Affymetrix UK Biobank Axiom array (67%) (Santa Clara, CA, USA) with the rest of the sample genotyped using the Affymetrix UK BiLEVE Axiom array. Quality control, phasing and imputation are described in detail elsewhere (http://biobank.ctsu.ox.ac.uk/crystal/refer.cgi?id = 155583), (http://biobank.ctsu.ox.ac.uk/crystal/refer.cgi?id = 155580), (http://biobank.ctsu.ox.ac.uk/crystal/refer.cgi?id = 157020). Briefly, phasing was performed using a modified version of the SHAPEIT2 algorithm^26^ with a combined panel of the UK10K and 1000 Genomes Phase 3 reference panels used for imputation^27^. Individuals were removed from the present study based on non-British ancestry (within those who self-identified as being British, principal component analysis was used to remove outliers, n = 32 484), high missingness (n = 0), relatedness (n = 7,948), QC failure in UK Bileve (n = 187), and gender mismatch (n = 0). For the GWAS analysis we used hard-called genotypes with an imputation info score ≥ 0.9, MAF ≥ 0.1% and HWE p-value ≤ 1 x 10^−6^ (n SNPs =12,489,782). SNPs were then filtered to only include those where >80% of the sample had a hard-called genotype.

*GS:* GS samples were genotyped using the Illumina Human

OmniExpressExome-8v1.0 Bead Chip and Infinum chemistry and processed using the Illumina Genome Studio Analysis software v2011. (Illumina, San Diego, CA). Quality control was performed to remove SNPs with <98% call rate, individuals with a genotyping rate <98%, and SNPs with a HWE p-value ≤ 1 x 10^−6^ and a MAF ≥ 1%. This left 561,125 SNPs available for analyses. More detail on blood collection and DNA extraction are provided elsewhere^24^.

### Genome-wide association analyses (GWAS)

In the UK Biobank, quality control was performed on the alcohol consumption phenotype to remove extreme values. Weekly intake values > 5 standard deviations from the mean were set to missing. For males this was for values > 102 units per week and for females for values > 89 units per week. As the mean alcohol intake in males was significantly higher than in females, we created the alcohol consumption phenotype by regressing age and weight in kilograms onto weekly units of alcohol consumed in males and females separately. We then took the residuals from these regressions and pooled the male and female residuals together to create the alcohol consumption phenotype. A male only and female only GWAS was also performed.

GWAS was performed using PLINK^28^ testing for associations between SNPs and alcohol consumption with location of UKB assessment center, genotyping batch and 15 principal components included as covariates. In order to distinguish independent GWAS signals, SNPs with an association p-value ≤ 1 x 10^−4^ were subjected to LD-based clumping was performed in PLINK^28^, using an LD r^2^ cutoff of 0.2 and a 500kb sliding window. Conditional SNP analyses were performed using PLINK for the 4 SNPs on chromosome 4q found to be associated with alcohol consumption. Each SNP was re-tested for association using the same method as described for the GWAS but using one of the three other SNPs as a covariate.

### Heritability and genetic correlation analyses

Univariate GCTA-GREML analyses were performed to estimate the SNP based heritability of alcohol consumption in the whole UK Biobank sample and then for males and females separately. n genetic relationship matrix was created for an unrelated subsample using a cut-off of 0.025 (N=89,175) and used to estimate heritability^29-31^. The genetic correlation between male and female alcohol consumption was estimated using LD score regression following the pipeline designed by Bulik-Sullivan and colleagues^32^. This method exploits the correlational structure of SNPs across the genome and uses test statistics provided from GWAS summary estimates to calculate the genetic correlations between traits^33^. FDR was used to correct p-values for multiple testing. We supplied the GWAS summary statistics from the GWAS of male and female alcohol consumption. Using our GWAS summary statistics, the genetic overlap between alcohol consumption and over 200 other disease traits was assessed using LD score regression^32^ implemented in the online software LD Hub (http://ldsc.broadinstitute.org/)^34^.

### Polygenic risk score analyses

Polygenic risk scores (PRS) were created in GS using raw genotype data using the software PRsice^35^ using the GWAS summary statistics from the UK Biobank GWAS of alcohol consumption. PRS were created using p-value thresholds ranging from 0.01 to 0.5 in increments of 0.01 using LD pruning parameters of r^2^ = 0.1 over 250kb windows. The PRS p-value threshold found to explain most of the variance in alcohol consumption in GS was at 0.25 and so this PRS was used to test for association in subsequent analyses. Association between alcohol PRS and traits of interest was performed in AS-Reml-R and an inverse relationship matrix created from the pedigree information in GS was used to control for relatedness in the sample. Four principal components were fit as fixed-effect covariates to control for population stratification. All traits and PRS were scaled to have a mean of 0 and a standard deviation of 1 such that the betas reported are standardized. The variance explained by PRS was calculated by multiplying the PRS by its regression co-efficient. This value was divided by the variance of the phenotype analysed to give a coefficient of determination between 0 and 1^36^. Bonferroni correction for PRS analyses required a p-value of 0.01 [0.05/5] as the threshold for statistical significance.

### eQTL and gene-based analyses

SNPs which were significantly associated with alcohol consumption (p< 5 x 10^−8^) in the GWAS were assessed to determine whether they were potential eQTLs using the Genotype Tissue Expression Portal (GTEx) (http://www.gtexportal.org). GTEx uses gene expression data from multiple human tissues linked to genotype data to provide information on eQTLs. Gene based analyses were performed using MAGMA^37^, derived from SNP summary data from the GWAS of alcohol consumption in this sample. The 1000 genomes European reference panel (phase1, release 3) was used to account for LD in the sample. SNPs were mapped to genes using the NCBI 37.3 gene locations and Entrez gene IDs. This resulted in 18,024 genes available for analysis which after Bonferroni correction (0.05/18,024) gave a threshold for statistical significance at 2.8 x 10^−6^.

## Results

The mean self-reported alcohol consumption in the 112,117 UK Biobank individuals who contributed to the analysis in the present study was 15.13 units per week (S.D.=16.56). The mean age of participants was 59.6 (S.D.=7.95). The sample comprised 59,088 females (52.7%). Females had a significantly lower mean weekly alcohol intake compared to males (10.03 (S.D.=11.83) vs 20.81 (S.D.=19.04) units per week), p ≤ 2 x 10^−16^.

The SNP-based heritability of alcohol consumption in the total sample was estimated to be 0.13 (S.E.=0.006). The SNP heritability in males was estimated to be 0.16 (S.E.=0.01) and in females 0.13 (S.E.=0.01). The SNP heritability in males was significantly higher than in females (Z-score = 1.963, p-value=0.0496). The SNP-based genetic correlation between male and female alcohol consumption using LD score regression was 0.73 (S.E.=0.09, p-value = 1.37 x 10^−16^) suggesting that the genetic factors influencing alcohol consumption in males and females in this sample were largely overlapping.

The genome-wide association statistics deviated slightly from the null (λ_**GC**_=1.086) (Supplemental Figure 1), however the LD score regression intercept of 1.007 suggested that the inflation of the test statistic was due to polygenicity rather than any population stratification^33^. Genome-wide significant associations (P < 5 x 10^−8^) were obtained for 10 loci after clump-based pruning (Table 1 and Figure 1). These included 4 hits on chromosome 4q23 which span ~250kb and several alcohol dehydrogenase genes (*ADH1B, ADH1C, ADH5*) which have all previously been implicated in alcohol consumption^9, 12^ One association on 4q23 is at a novel locus, *DNAJB14.* These are all relatively rare variants with a MAF ranging from 0.005-0.01 with the minor allele associating with lower alcohol consumption (Table 1). Conditional SNP GWAS of SNPs on chromosome 4q suggests that there 2 completely independent hits in this region: rs145452708 in the *ADH1B/ADH1C* region and rs29001570 in *ADH5.* The SNP rs29001570 in *ADH5* remained significantly associated with alcohol consumption when any of the other 3 significant SNPs on 4q were used as a covariate in the GWAS (Supplementary Table 1). However, the other 3 SNPs either became non-significant or showed marked attenuation in the strength of the association with alcohol consumption when conditioned on one-another. Adjusting for top SNP rs145452708 in the *ADH1B/ADH1C* region resulted in the effects of rs144198753 (β=-0.005, p=0.28) and rs140280172 (β=-0.001, p=0.72) becoming non-significant. Two hits were identified on chromosome 4p14 in the *KLB* gene, which was previously associated with alcohol consumption in a large GWAS of >105,000 Europeans^14^. Novel hits included a SNP on chromosome 2p23.3 in the *GCKR* gene. The SNPs in *KLB* and *GCKR* were the most strongly associated common (MAF >5%) SNPs with MAFs of >0.28. Regional association plots for the common SNPs in *KLB* and *GCKR* are shown in Figure 2. Additional novel hits on chromosome 2p25.3 (*PXDN*), chromosome 3p12.1 *(CADM2)* and chromosome 18q21.33 (*TNFRS11A*) were also identified (Table 1). These loci will be reviewed in greater detail in the discussion. Regional association plots for all SNPs presented in Table 1 are shown in Supplementary Figures 2-9. A GWAS of alcohol consumption was performed excluding individuals who identified as never drinkers. The SNPs associated with alcohol consumption in current drinkers were the same as those reported in the total sample with the exception of rs439945 in the corticoptropin releasing hormone receptor 1 (*CRHR1*) gene (Supplementary Table 2 and Supplementary Figure 10). Sex-specific GWAS of alcohol consumption were also performed (Supplementary Figures 11 and 12). Only one novel locus was identified in males on chromosome 16 in a zinc-finger protein gene (*ZNF689*). rs548307545 was not significantly associated with alcohol consumption in females (β=-0.007, S.E.=0.004, p=0.12). All other loci identified in males or females had previously been identified at the level of genome-wide significance in the total sample (Supplementary Table 3).

**Table 1:** Ten loci reaching genome-wide significance for association with alcohol consumption in UKB after performing clump-based LD pruning. Genes are reported if SNPs located +/- 10kb of the locus. CHR= chromosome, POS= base pair position, Freq= A1 frequency

| SNP | CHR | POS | A1/A2 | Freq | $\beta$(se) | P | Genes |
| --- | --- | --- | --- | --- | --- | --- | --- |
| rs145452708 | 4 | 100248642 | C/G | 0.01 | -0.033 (0.003) | $8.93 \times 10^{-29}$ | *ADH1B/ADH1C* |
| rs144198753 | 4 | 99713350 | T/C | 0.008 | -0.029 (0.003) | $7.57 \times 10^{-23}$ | - |
| rs28712821 | 4 | 39413780 | G/A | 0.39 | -0.027 (0.003) | $1.05 \times 10^{-18}$ | *KLB* |
| rs1260326 | 2 | 27730940 | T/C | 0.39 | -0.025 (0.003) | $4.67 \times 10^{-17}$ | *GCKR* |
| rs29001570 | 4 | 99994405 | C/T | 0.006 | -0.024 (0.003) | $2.46 \times 10^{-15}$ | *ADH5* |
| rs140280172 | 4 | 100832564 | A/C | 0.005 | -0.020 (0.003) | $4.63 \times 10^{-11}$ | *DNAJB14* |
| rs9991733 | 4 | 39420994 | G/A | 0.28 | 0.018 (0.003) | $6.18 \times 10^{-10}$ | *KLB* |
| rs9841829 | 3 | 85569361 | G/T | 0.23 | 0.017 (0.003) | $4.52 \times 10^{-9}$ | *CADM2* |
| rs187953385 | 2 | 1702871 | T/C | 0.001 | 0.017 (0.003) | $6.77 \times 10^{-9}$ | *PXDN* |
| rs4447524 | 18 | 60034276 | C/T | 0.15 | 0.017 (0.003) | $3.89 \times 10^{-8}$ | *TNFRSF11A* |

**Figure 1:**
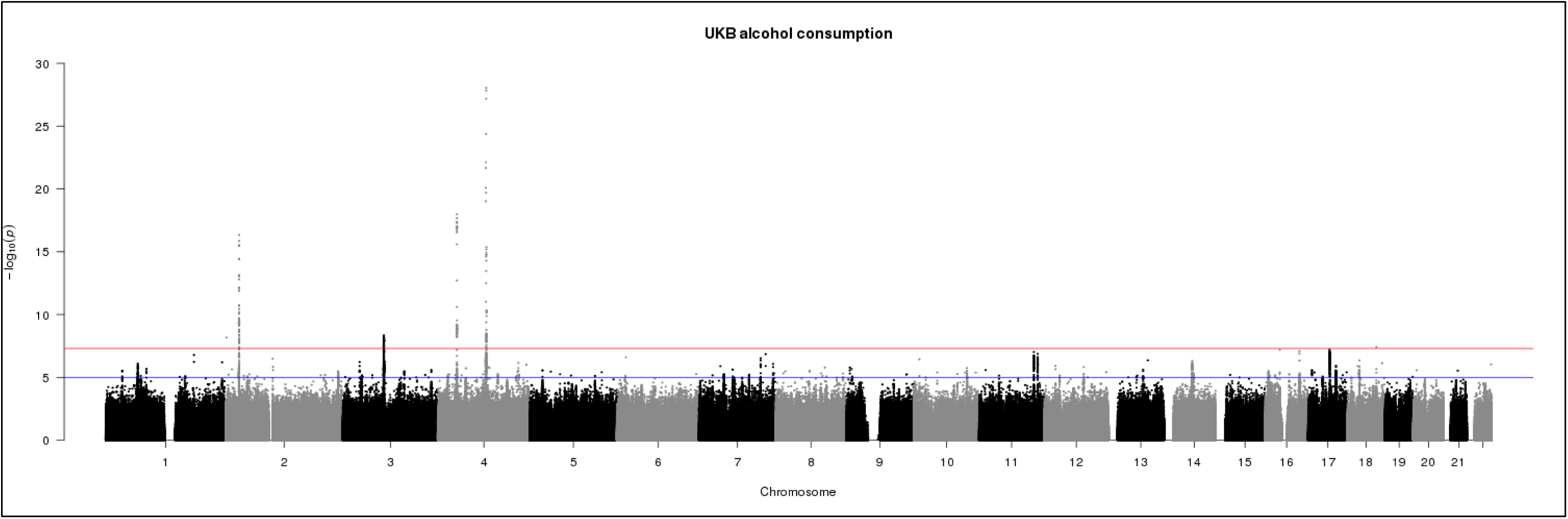
Manhattan plot of GWAS of alcohol consumption in UKBiobank (N=112,177). Red line indicates threshold for genome-wide significance (p ≤ 5 x 10^−8^) and the blue line for suggestive significance (p ≤ 5 x 10^−6^).

**Figure 2:**
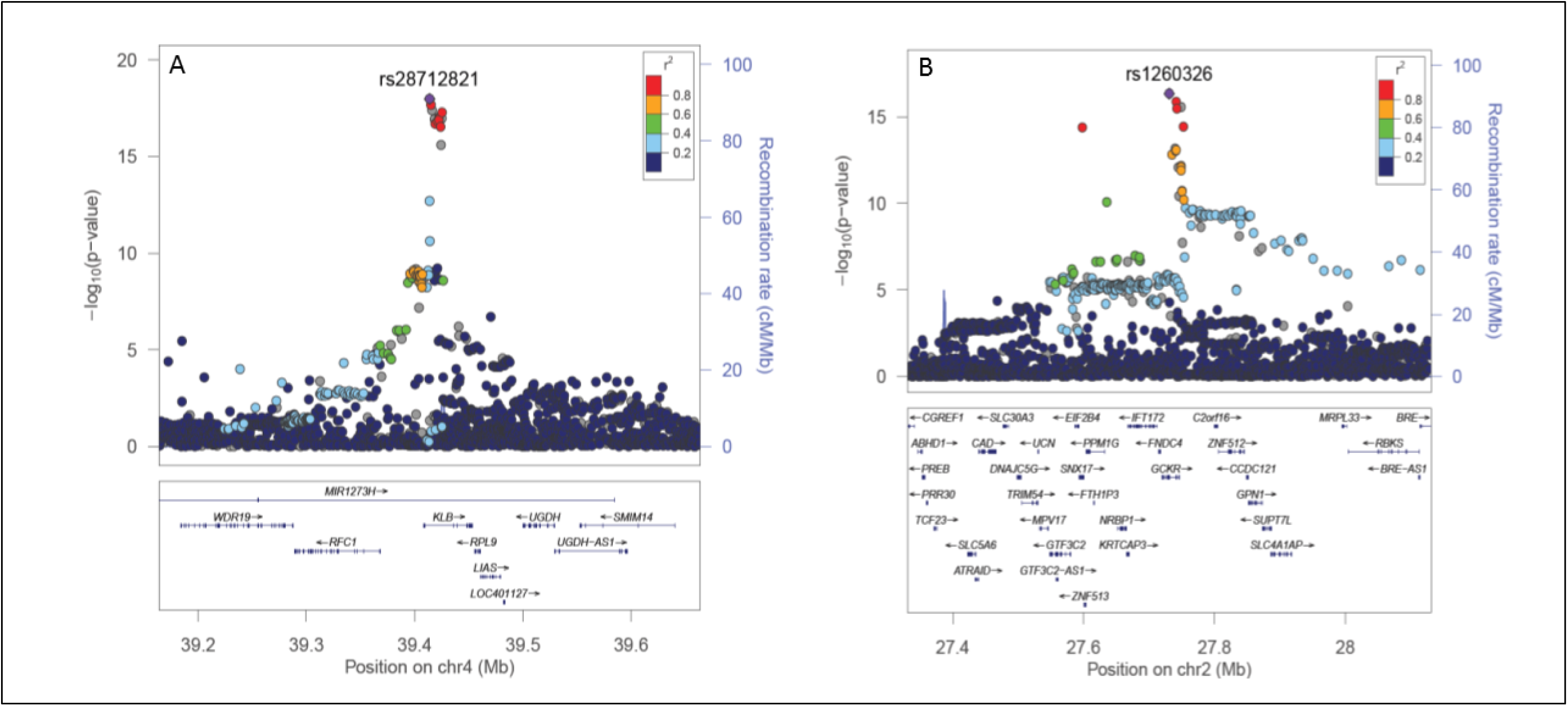
LD zoom plots of 2 common SNPs (MAF >5%) most significantly associated (P < 5 x 10^−8^) with alcohol consumption in UKBiobank. A: rs2872821 on chromosome 4 in *KLB* and B: rs1260326 on chromosome 2 located in *GKCR.*

The gene-based analyses found 36 genes to be significantly associated with alcohol consumption (Supplementary Table 4). The top hits were genes identified in the single SNP analyses (*KLB, CADM2, GCKR*) and genes in the 4q region were also found to be significantly associated with alcohol consumption at the gene-level (*ADH1C, C4orf17*). *CRHR1* was associated at the gene level and also associated with alcohol consumption at the SNP level when never drinkers were excluded. Amongst the 36 loci associated at the gene-level, genes of interest include the dopamine receptor D2 gene (*DRD2*) previously associated with addiction phenotypes^38^, autism susceptibility gene 2 (*AUTS2*) previously associated with alcohol consumption^13^, cAMP-specific 3’,5’-cyclic phosphodiesterase 4B (*PDE4B*) associated with alcohol preference in rodents^39, 40^, zinc-finger protein 512 (*ZNF512*) associated with oral cavity cancer^41^, and protein phosphatase 1G (*PPM1G*) found to be hyper-methylated in individuals with alcohol use disorders^42^.

Four SNPs located across *KLB, CADM2* and *GCKR* were found to be eQTLs according to the GTEx database (Supplementary Table 5). Notably, rs28712821 in *KLB* was found to be an expression QTL for *RCF1* and *RPL9* in the cerebellum. rs9841829 in *CADM2* is associated with expression of *CADM2* levels in lung and adipose tissue.

The genetic correlations between alcohol consumption and 219 other health and behavioural traits were calculated using GWAS summary statistics and LD score regression^32^ implemented in the online software LD Hub^34^. After correction for multiple testing 10 traits had a significant genetic correlation with alcohol consumption (Figure 3). Smoking status had the strongest positive genetic correlation with alcohol consumption (rG=0.49, S.E.=0.06, p=1 x10^−14^) followed by lung cancer (rG=0.25, S.E.=0.07, p=6 x10^−4^) and HDL cholesterol levels (rG=0.21, S.E.=0.05, p=2 x10^−5^). Significant negative genetic correlations were observed for childhood height (rG=-0.21, S.E.=0.06, p=4 x10^−4^) and a range of other anthropometric traits pertaining to weight and obesity, such as obesity class 2 (rG=-0.20, S.E.=0.06, p=4 x10^−4^) which categorizes severely obese individuals with a BMI ranging from 35.0 to 39.9. Genetic correlations were also calculated using the sex-specific GWAS summary statistics (Supplementary Figures 13 and 14). Male alcohol consumption showed the same significant genetic correlations that were identified in the total sample. However, female alcohol consumption additionally found college completion (rG=0.22, S.E.=0.07, p=8 x10^−4^) and years of schooling (rG=0.15, S.E.=0.04, p=3 x10^−4^) to be positively genetically correlated with alcohol consumption. No significant genetic correlations between college completion (rG=-0.05, S.E.=0.05, p=0.34) or years of schooling (rG=-0.04, S.E.=0.04, p=0.30) were observed with male alcohol consumption. Several metabolic traits were found to have negative genetic correlations with female alcohol consumption including LDL cholesterol (rG=-0.25, S.E.=0.07, p=5 x10^−4^) and triglycerides (rG=- 0.23, S.E.=0.07, p=3 x10^−2^).

**Figure 3:**
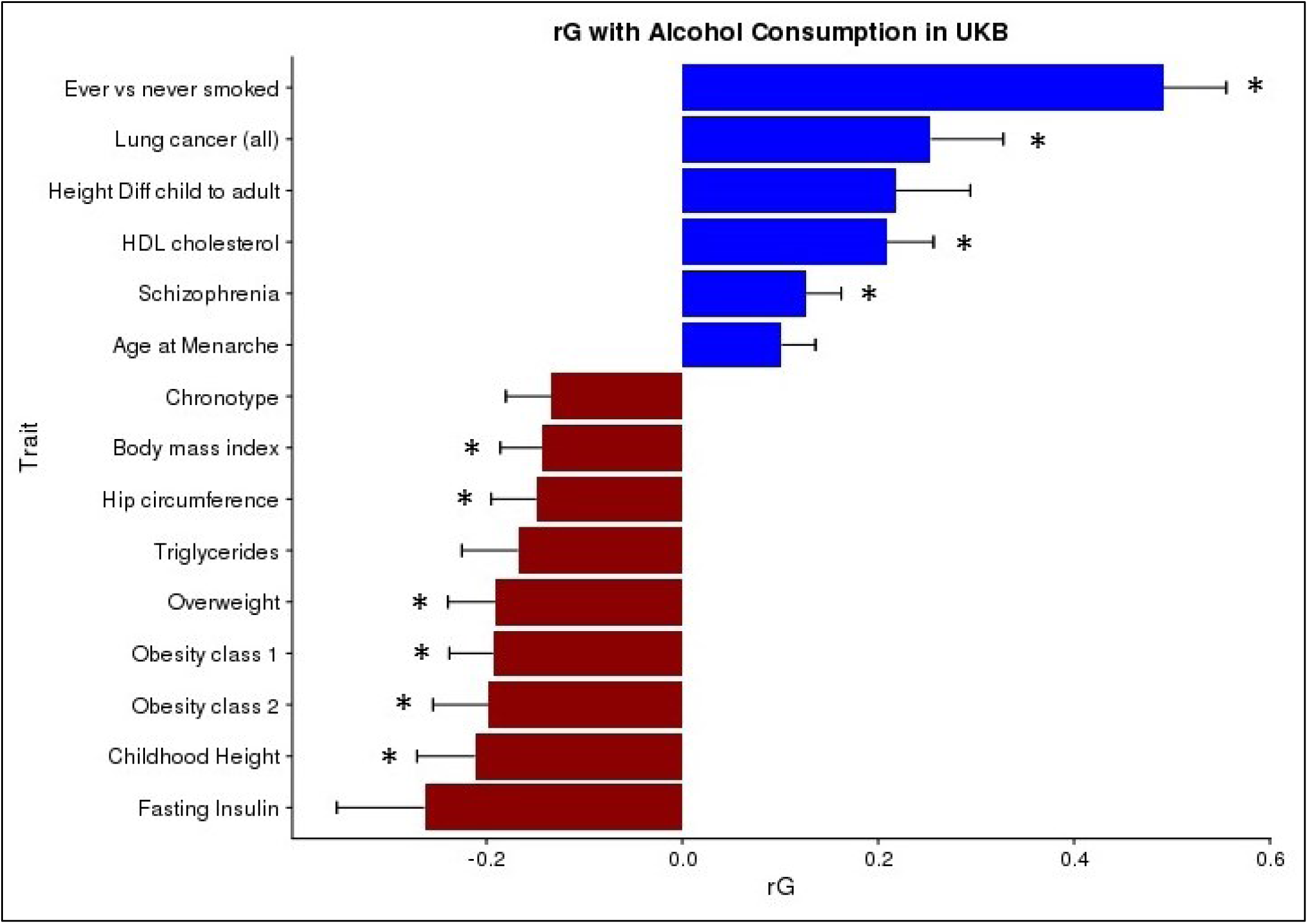
Genetic correlation between Alcohol Consumption in UKB and other traits using LD score regression implemented in LDHub. Childhood Height = Females at age 10 and males at age 12, Height Diff child to adult = Difference in height between childhood and adulthood; age 8. * = statistically significant after FDR correction for multiple testing. Traits presented that were not significant after FDR correction have an FDR p-value < 0.06.

Using the UK Biobank’s GWAS summary statistics PRS for alcohol consumption were calculated in GS to determine the amount of variance that could be explained using a risk-score approach. PRS for alcohol consumption in GS were found to be positively associated with alcohol consumption (β=0.07, p=6.8 x 10^−21^) however only 0.5% of the variance in alcohol consumption was explained (Table 2). PRS were also tested for association with traits found to have a significant genetic correlation with alcohol consumption in the LD score regression analyses. BMI, weight, hip circumference, smoking status and HDL cholesterol levels were all available in the GS cohort. Significant associations between alcohol PRS and smoking were detected after adjustment for multiple testing (β=-0.143, p=0.00013), and nominally significant associations with HDL cholesterol (β=0.016, p=0.02) (Table 2); however, no significant associations were found between PRS scores and any of the anthropometric traits.

**Table 2:** PGRS analyses: PGRS for alcohol consumption in Generation Scotland created using summary statistics of alcohol consumption GWAS in UKB. Association analyses performed in AS-Reml-R using pedigree information to control for relatedness. Units per week phenotype for current drinkers only. All models adjusted for age, sex and 4 MDS components to control for population stratification. Bold highlighted p-values are statistically significant after correction for multiple testing (p ≤ 0.01).

| **Trait (n)** | **$\beta$(se)** | **Z Ratio** | **r<sup>2</sup>** | **p-value** |
| --- | --- | --- | --- | --- |
| Units per week (17,461) | 0.070 (0.007) | 9.39 | 0.5% | **6.8 x 10<sup>-21</sup>** |
| Ever smoke (19,289) | -0.143 (0.004) | -3.82 | 0.08% | **0.00013** |
| BMI (19,771) | -0.007 (0.007) | -0.98 | 0.005% | 0.33 |
| Hips (19,58) | 0.0006 (0.008) | 0.08 | 0.0004% | 0.94 |
| HDL Cholesterol (19,108) | 0.016 (0.007) | 2.28 | 0.03% | 0.02 |

## Discussion

In the present study we identify 8 independent loci that are associated with alcohol consumption. Two of these loci are located amongst a cluster of alcohol metabolism genes on chromosome 4q23 (*ADH1B/ADH1C* and *ADH5*) and have been previously identified as risk loci for alcohol related phenotypes^9, 12^ Two SNPs in *KLB* were also associated with alcohol consumption and this locus has previously been identified in a large meta-analysis of alcohol consumption in Europeans^14^. The remaining 4 hits are novel loci (*GCKR, CADM2, TNFRSF11A*, and *PXDN*) in the context of alcohol consumption and as such this study presents a novel contribution to the genetics of alcohol consumption.

Identifying the causal variants located on 4q23 will prove to be challenging due to the low MAF of the variants associated with alcohol consumption in this region. Previous studies have shown that rs1229984 in *ADH1B,* which is associated with rapid alcohol metabolism, is strongly protective against high alcohol consumption^10^. It is possible that the low frequency variants identified in 4q23 are tagging rs1229984. However, rs1229984 deviated from HWE in the UK Biobank sample (HWE p=1.5 x 10^−78^) and therefore was not analysed as part of the GWAS. rs29001570 in *ADH5* clearly represents an independent locus as the r^2^ with the 3 other SNPs in 4q23 is < 0.01. This is the first GWAS of alcohol consumption to detect genome-wide significant association with *ADH5,* although a GWAS of alcohol dependence found suggestive evidence of association with *PDLIM5,* which is adjacent to ADH5^12^. The 3 other 4q23 SNPs are in moderate LD (r^2^ = 0.23-0.68) (Supplementary Table 6) and conditional SNP analyses suggest that there is likely to be only a single independent variant at this locus.

The gene encoding β-Klotho (*KLB*) has recently been associated with alcohol consumption in a large meta-analysis of >105,000 Europeans. Schumann and colleagues found that brain-specific *klb*-knockout mice have increased alcohol preference. They also found that β-Klotho is a receptor for the liver expressed hormone FGF21 which acts on the brain and inhibits alcohol consumption in mice^14^. We are not aware of any sample overlap between UKB and the sample used in the study by Schumann and colleagues. Generation Scotland individuals did contribute to the UKB cohort sample however GS individuals and relatives of GS participants were removed from the UKB sample prior to GWAS analysis. The study by Schumann and colleagues identified rs11940694 as associated with alcohol consumption which is in complete LD with the SNP in *KLB* identified in the present study, rs28712821 (r^2^=1, D’=1). We therefore provide evidence of replication of the association of this variant with alcohol consumption. An additional SNP (rs9991733) in *KLB* was associated with alcohol consumption in UKB however the minor allele of this SNP conferred risk for high consumption rather than protection. This SNP is in low LD with rs28712821 (r^2^=0.21, D’=0.87) demonstrating that there are multiple variants in *KLB* that are associated with alcohol consumption.

Novel loci found to be associated with alcohol consumption in this study include *GCKR, CADM2, TNFRSF11A* and *PXDN.* With the exception of *CADM2,* these genes encode proteins that have roles in the liver, are implicated in liver disease or are associated with alcoholic liver pathology. *GCKR* encodes the glucokinase regulatory protein which is produced by hepatocytes and is responsible for phosphorylation of glucose in the liver. rs1260326 in *GCKR* is a coding mis-sense SNP and this variant has been associated with over 25 metabolic traits including type II diabetes, fasting insulin levels and total cholesterol levels^43^. *TNFRSF11A* encodes the receptor activator for NF-κB (RANK). The main RANK ligand is RANKL and together with its decoy receptor osteoprotegerin (OPG) is involved in osteoclast formation and immune function^44^. The RANK/RANKL/OPG system has been shown to be upregulated in alcoholic liver cirrhosis^45, 46^ and correlated with disease severity in an auto-immune liver disease, primary biliary cholangitis^44^. *PXDN* encodes the peroxidasin homolog which is involved in extracellular matrix formation. A *de novo* variant in *PXDN* was found in an individual with congenital dilatation of the bile-duct, a liver disorder resulting in bile retention^47^. *CADM2* is a brain expressed gene encoding cell adhesion molecule 2 which has previously been associated with processing speed and educational attainment in the UKB sample and processing speed in the CHARGE consortium cohort^48, 49^.

Several additional genes associated with alcohol consumption were identified through MAGMA gene-based analyses, including the gene *CRHR1.* This gene encodes the corticotrophin-releasing hormone receptor 1 which facilitates hypothalamic-pituitary-adrenal (HPA) axis signalling and as such mediates the stress response. CRH signalling becomes important as substance use escalates to substance addiction by contributing to anxiety states, reward deficits and stress-induced re-instatement of substance use (reviewed in, Zorilla et al.^50^). Polymorphisms in *CRHR1* have been linked to adolescent alcohol consumption and levels of consumption amongst alcohol dependent individuals^51^. Furthermore, several studies have found that SNPs in *CRHR1* interact with various forms of life stress to influence alcohol consumption levels^52-54^.

*AUTS2* was also associated with alcohol consumption at the gene-level and a SNP in this gene was previously found to be associated with alcohol consumption in a GWAS of 47,501 Europeans^13^. *PDE4B* was associated at the gene level and encodes a protein which regulates alcohol induced neuro-inflammation in the CNS^55^. Animal models have shown that inhibition of PDE4B leads to reduced alcohol intake in both mice and rats^39, 40^. A GWAS found *ZNF512* to be associated with oral cavity cancer^41^. The gene-based analysis in the present study found this gene to also be associated with alcohol consumption. This is notable considering that alcohol consumption is a leading cause of oral cavity cancer, suggesting that the link between *ZNF512* and alcohol consumption may underlie its association with oral cancer.

Using LD score regression, significant positive genetic correlations between alcohol consumption and smoking and lung cancer were identified. The overlap between alcohol consumption and smoking is well documented and other studies have shown polygenic overlap between weekly alcohol intake and the number of cigarettes smoked (rG=0.44)^20^. It is likely that the relationship between alcohol consumption and smoking underlies the positive genetic correlation with lung cancer detected in the present study. A positive genetic correlation between alcohol consumption and schizophrenia was also detected (rG=0.13). Alcohol abuse has been shown to increase the risk for schizophrenia^56^; however, a twin study of schizophrenia and alcoholism found no evidence of genetic overlap between the traits^57^. This is the first study, to our knowledge, to report a genetic correlation between schizophrenia and alcohol consumption. A positive genetic correlation between HDL cholesterol and alcohol consumption was also found. Increased alcohol consumption is associated with increased HDL levels^58^ and a Mendelian randomization study of alcohol consumption and lipid profiles found a causal effect of alcohol on increased HDL levels in the low to moderate intake range^59^. rs1260326 in *GCKR* has previously been associated with HDL levels^60^ and this was one of the most significant associations with alcohol consumption in the present study. Furthermore, alcohol PRS in GS were nominally associated with increased HDL. Our findings provide further support for the positive relationship between alcohol and HDL levels.

Negative genetic correlations were observed for several anthropometric traits including BMI and obesity, such that genetic variants which increased risk for alcohol consumption decrease the risk for overweight phenotypes. Previous studies have shown both negative and positive relationships between alcohol and BMI^61, 62^ and a SNP in the *FTO* gene coding region which increases risk for obesity also increases risk for alcohol dependence^63^. We report a negative genetic correlation between obesity related traits and alcohol consumption; albeit this was not replicated using a PRS approach in GS possibly due to the smaller sample size of GS and the limited power of PRS approaches. A negative genetic correlation between alcohol consumption and childhood height was observed and a nominally significant positive genetic correlation with height difference between childhood height at age 8 and at adulthood. Alcohol consumption has been associated with delayed puberty^64, 65^ and this is now known to occur in part through alcohol induced inhibition of the gonadotropin releasing hormone (GnRH) ^66-69^ Alcohol blocks GnRH signalling by inhibiting transforming growth factor beta 1 (TGFβ1) signalling^70^. Interestingly, we find association of *TGFB1* in our gene-level analyses demonstrating that polymorphisms in this gene associate with adult alcohol consumption. Future work may include identifying whether specific genetic variants in *TGFB1* associate with differential effects of alcohol on pubertal processes in humans.

This investigation has several advantages compared to previous studies. We studied a single large sample of ancestrally similar individuals from a relatively narrow age range, all of whom were residing in the United Kingdom at the time of interview. Using a sample where all individuals are exposed to similar social norms is advantageous as cultural influences are an important influence on alcohol intake patterns^71^. Furthermore, all data was subject to the same quality control procedures and these factors will have contributed to our ability to detect 8 independent genome-wide significant loci when similar sized alcohol consumption GWAS did not^14^. The main limitation of this study is the reliance on self-reported alcohol consumption and the reliability of this measure. However, the UKB touchscreen interview used to ascertain alcohol intake used pictures to represent different drink sizes which may have increased reliability. Finally, the results presented here may only be applicable to middle to older aged white British individuals. Considering that genetic influences on alcohol consumption change across the life course^72^ it is important to note that the variants identified in this study may have limited relevance outside of this demographic.

This study presents the largest single GWAS of alcohol consumption to date and identifies 8 genetic loci, at least 4 of which are novel. The SNP heritability of alcohol consumption is described for the first time with 13% of the variance in alcohol consumption attributable to genetic factors in this sample. Many of the genes identified in this study are expressed in the liver and are associated with liver pathology suggesting that alcohol metabolism is an important driver of differences in consumption. Finally, the genetic overlap with alcohol consumption and novel traits such as schizophrenia are reported for the first time. The genetic overlap between alcohol consumption and HDL cholesterol is confirmed at the level of single SNP association and overlapping polygenic architecture. Future studies should focus on characterizing the biological role of these loci in alcohol consumption and investigating the potential causal relationships between alcohol use and health related traits.

## Acknowledgements

This research has been conducted using the UK Biobank Resource – application number 4844.

We are grateful to the families who took part in GS, the GPs and Scottish School of Primary Care for their help in recruiting them, and the whole GS team, which includes academic researchers, clinic staff, laboratory technicians, clerical workers, IT staff, statisticians and research managers. Generation Scotland received core support from the Chief Scientist Office of the Scottish Government Health Directorates [CZD/16/6] and the Scottish Funding Council [HR03006]. Genotyping of the GS samples was carried out by the Genetics Core Laboratory at the Wellcome Trust Clinical Research Facility, Edinburgh, Scotland and was funded by the Medical Research Council UK and the Wellcome Trust (Wellcome Trust Strategic Award “STratifying Resilience and Depression Longitudinally” (STRADL) Reference 104036/Z/14/Z). We acknowledge with gratitude the financial support received for this work from the Dr Mortimer and Theresa Sackler Foundation. PT, DJP, IJD, and AMM are members of The University of Edinburgh Centre for Cognitive Ageing and Cognitive Epidemiology, part of the cross council Lifelong Health and Wellbeing Initiative (MR/K026992/1). Funding from the Biotechnology and Biological Sciences Research Council and Medical Research Council is gratefully acknowledged.

